# Nmur1 and Cckar fail to support functional genetic access in adult dopamine neurons and challenge GPCR atlas assignments

**DOI:** 10.64898/2026.05.11.724447

**Authors:** Moueez Shah, Renqi Wu, Qiao Ye, Raluca Bugescu, Andrew P. Villa, Justine Wong, Fernando Garcia, Zhiqun Tan, Xiangmin Xu, Gina M. Leinninger, Andrew D. Steele

## Abstract

Apuschkin et al. (2024) proposed a GPCR-based transcriptomic atlas for midbrain dopamine (DA) neuron subpopulations, including candidates such as *Nmur1, Cckar*, and *Ffar4*. To guide genetic targeting, these markers must reflect functional expression in adult DA neurons. Using *in situ* hybridization, Cre-dependent reporter lines, and both intracranial and systemic viral approaches, we find no evidence of adult *Nmur1*-mediated recombination in DA neurons, while *Cckar*-driven recombination is consistent with developmental expression only. Notably, *Ffar4* expression overlaps extensively with *Ntsr1* midbrain populations, indicating that it does not define a distinct DA neuron class. Furthermore, analysis of independent spatial transcriptomic datasets together with our MERFISH data shows that many proposed GPCR markers are not detectably expressed in adult DA neurons. These findings demonstrate that transcriptomic enrichment does not always yield reliable adult markers and highlight the need for functional validation prior to use in circuit targeting.

## Introduction

Recent advances in spatial, single-cell and single-nucleus transcriptomics have enabled increasingly detailed classification of midbrain dopamine neuron subpopulations.^1–3^ These approaches have identified candidate molecular markers, including G protein-coupled receptors (GPCRs), that are proposed to define functionally distinct dopamine (DA) neuron groups.^4^ Because GPCRs are both pharmacologically tractable and genetically accessible through Cre mouse lines or ‘enhancer’ adeno-associated viruses (AAV), such datasets are increasingly used to guide experimental targeting strategies in studies of DA function within the *substantia nigra* (SN) and ventral tegmental area (VTA).

However, transcriptomic enrichment does not necessarily translate into reliable markers of adult neuronal identity. Low-level expression, transient developmental expression, and technical limitations of detection can all lead to misassignment of candidate markers when these genes are used to drive Cre-dependent genetic manipulations.^2,5^ While recent GPCR atlas approaches of Apuschkin and colleagues (2024) nominate dozens of candidate receptors as population markers for DA neurons, experimental validation has been limited, with functional characterization focused on a single GPCR, *Free fatty acid receptor 4* (*Ffar4*), despite the breadth of candidates identified.^4^ Furthermore, *Ntsr1* and *Crhr1*, both well-established midbrain DA GPCR markers were either absent or showed very restricted expression in their analysis. For example, Apuschkin and colleagues report that only 6% of midbrain DA neurons express *Ntsr1*^4^ whereas other studies have demonstrated broad expression of *Ntsr1* in midbrain DA neurons.^6,7^ This gap raises important questions regarding the extent to which this dataset can be directly translated into genetic targeting strategies.

Because Cre driver lines are readily available for many GPCRs, we focused our analysis on two candidates from the GPCR atlas of midbrain DA neurons: *Neuromedin U Receptor 1* (*Nmur1*) and *Cholecystokinin A Receptor* (*Cckar*).^4^ Here, we evaluated their expression in adult mouse midbrain using complementary approaches, including a Cre-dependent reporter line, directly injected and systemically injected reporter AAVs, *in situ* hybridization, and conditional genetic manipulations. We find that *Nmur1* is not detectably expressed in adult DA neurons and does not drive functional recombination in this population. In contrast, *Cckar*-driven recombination is consistent with developmental expression only. These findings indicate that *Nmur1* and *Cckar* do not reliably define adult DA neuron subpopulations. Consistent with these findings, independent spatial transcriptomic datasets provide limited support for many proposed GPCR markers in adult dopaminergic neurons. Analysis of publicly available spatial transcriptomic resources, including the Allen Brain Cell (ABC) Atlas and Broad Institute datasets, together with our own data from multiplexed error-robust fluorescence in situ hybridization (MERFISH) experiments, reveals limited or absent expression of many candidate GPCRs within *Th*^*+*^ midbrain dopaminergic populations. Together, our findings suggest that some GPCRs identified in the atlas may reflect sparse, transient, developmentally restricted, or false positives rather than stable adult dopaminergic population markers. These findings underscore the importance of validating GPCR expression prior to their use in functional studies of dopamine circuits and behavior.

## Results

To examine whether *Nmur1* and *Cckar* are expressed within DA midbrain populations, we crossed *Nmur1*^*iCre*^ and *Cckar*^*iCre*^ mice to an *Ai6* (*lox-stop-lox ZsGreen1*) reporter line^8^ to assess Cre-mediated recombination.^8–10^ In *Nmur1*^*iCre*^;*Ai6* mice we observed sparse ZsGreen1 reporter expression in colocalizing with tyrosine hydroxylase (TH): approximately 2% of TH^+^ neurons co-labeled in the SN and 5% in the VTA (**Figure 1A-B**). In contrast, *Cckar*^*iCre*^*;Ai6* mice displayed more ZsGreen1^+^cells in both the SN and VTA, with approximately 5% of TH^+^ neurons labeled in the SN and 25% in the VTA (**Figure 1 C-D**).

**Figure 1.**
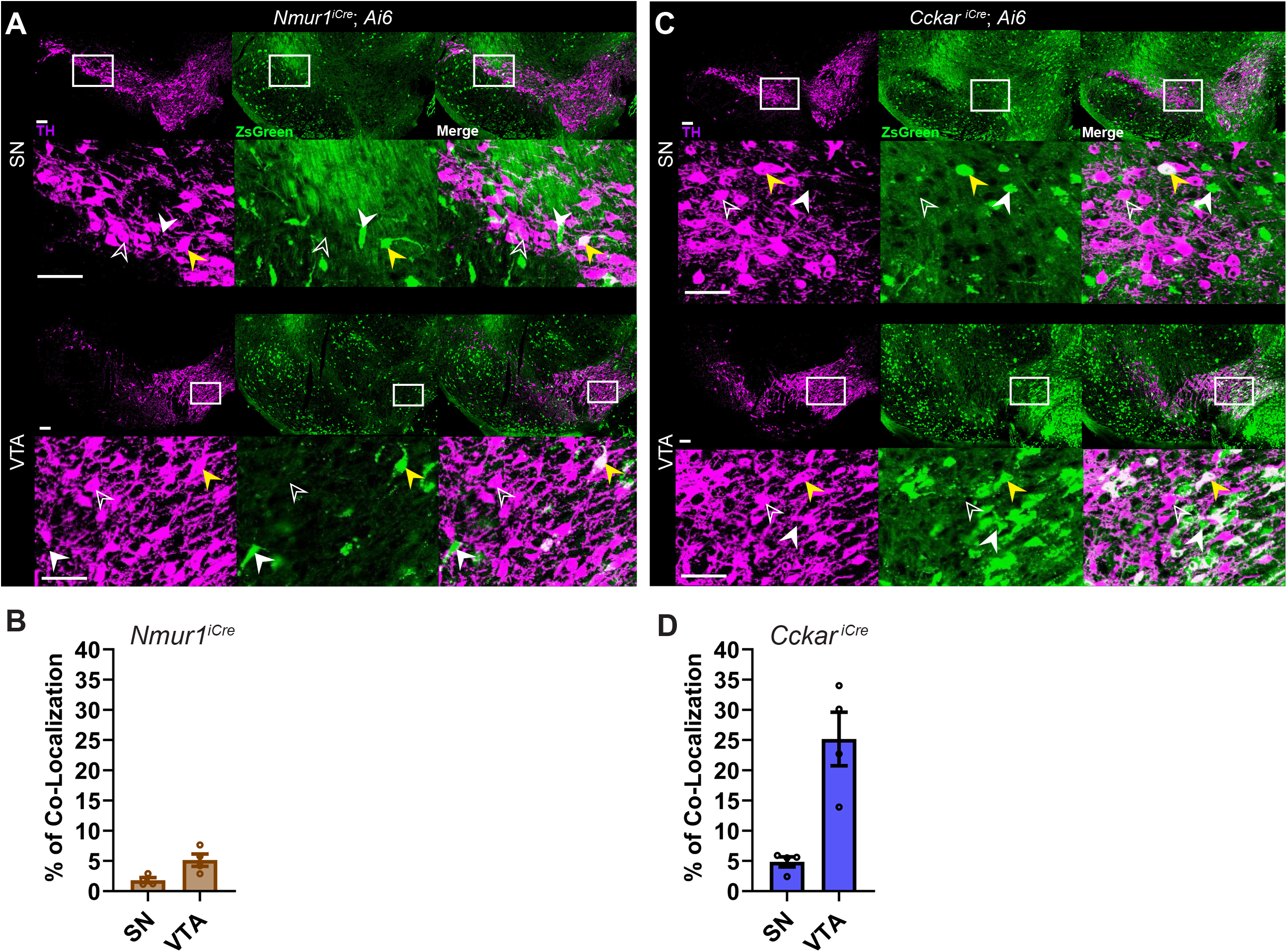
Sparse and differential labeling of midbrain dopamine neurons in *Nmur1*^*iCre*^*;Ai6* and *Cckar*^*iCre*^ *;Ai6* mice. (A, C) Representative images of ZsGreen reporter expression in the substantia nigra (SN) and ventral tegmental area (VTA) from *Nmur1*^*iCre*^*;Ai6* and *Cckar*^*iCre*^*;Ai6* mice. TH immunostaining (magenta) and ZsGreen expression (green). Merged image showing colocalization of Zsgreen and TH (white). Insets show higher-magnification views of boxed regions. Yellow arrowheads indicate TH^+^/ZsGreen colocalization. Open arrowheads indicate TH^+^ cells. White arrowheads indicate ZsGreen reporter. (B, D) Quantification of co-localization between reporter-positive cells and TH^+^ neurons in SN and VTA. Data are shown as mean ± SEM. Sample sizes: n = 4. Scale bars = 100 *µ*m.

To determine whether low-level gene expression might nonetheless be sufficient to drive functional recombination, we employed a Cre-lox-based strategy to selectively delete *Th* in a Cre-defined population. With this approach, we generated two conditional knockout (cKO) lines, *Nmur1*^*iCre*^*;Th*^*Flox*^ and *Cckar*^*iCre*^*;Th*^*Flox*^. In contrast to what would be expected for a *bona fide* DA neuronal marker, conditional deletion of *Th* in *Nmur1*^*iCre*^*;Th*^*Flox*^ mice failed to alter TH+ neuron number or distribution in a meaningful manner (SN: p = 0.5360, VTA: p = 0.2045, two-tailed unpaired t test; **Figure 2A-B**). In contrast, *Cckar*^*iCre*^*;Th*^*Flox*^ mice displayed a more pronounced reduction in TH^+^ neurons relative to controls, with the decrease apparent in both VTA and SN (SN: p = 0.0028, VTA: p = 0.0107, two-tailed unpaired t test; **Figure 2C-D**). This corresponds to an approximate 35% reduction in SN and 25% in VTA.

**Figure 2.**
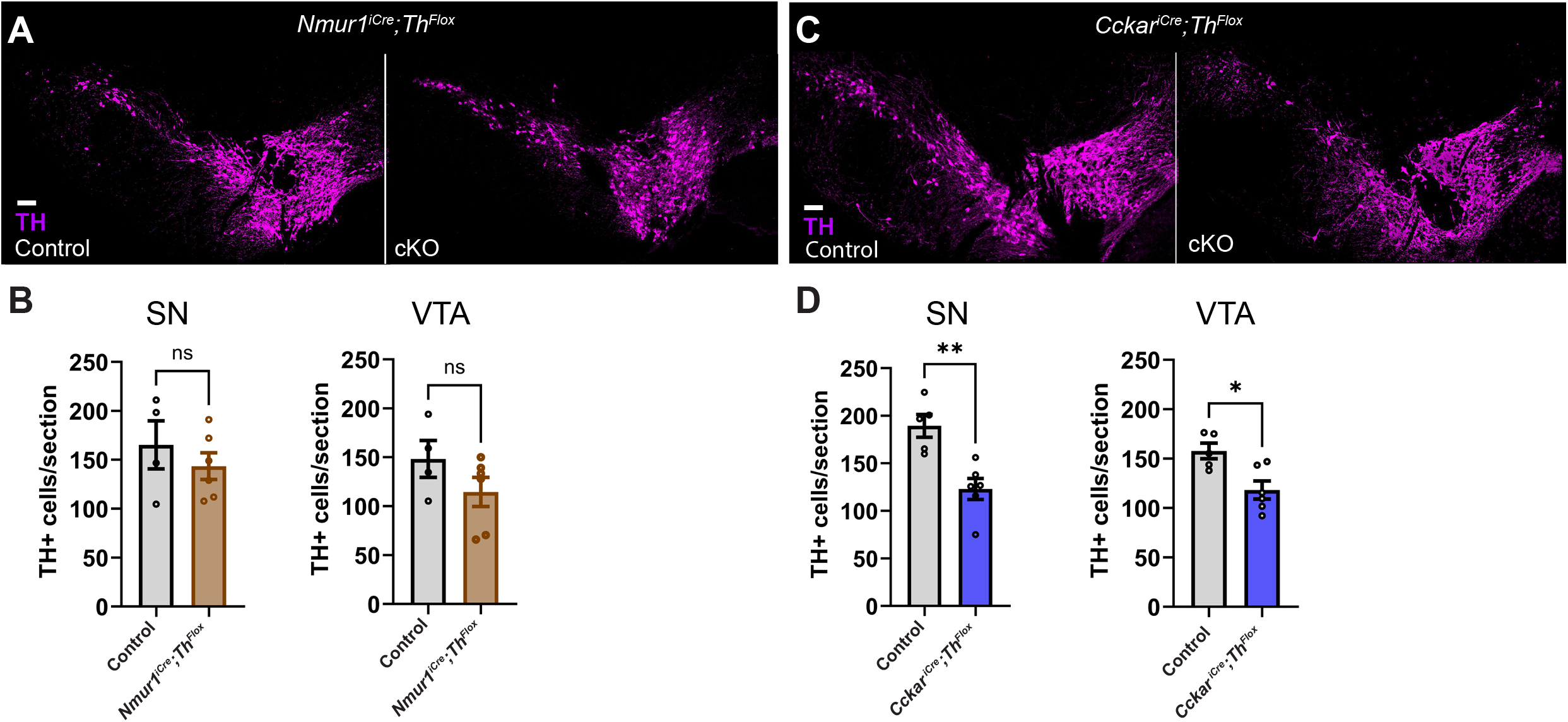
Conditional deletion of Th reveals limited recombination in *Nmur1* neurons and substantial targeting in *Cckar* populations. (A) Representative images of TH (magenta) immunolabeling in the SN and VTA from control and *Nmur1*^*iCre*^*;Th*^*Flox*^ mice. (B) Quantification of TH+ neurons in the SN and VTA of control and *Nmur1*^*iCre*^*;Th*^*Flox*^ mice (C) Representative images of TH immunolabeling in control and *Cckari*^*Cre*^*;Th*^*Flox*^ mice. (D) Quantification of TH+ cells/section ± SEM in the SN and VTA of control and *Cckari*^*Cre*^*;Th*^*Flox*^ mice. Data are shown as mean ± SEM. Statistically significant differences are indicated as follows: ns, non-significant; *p < 0.05; **p < 0.01. Sample sizes: n = 5 per group. Scale bars = 100 *µ*m.

Next, to assess adult Cre activity using viral reporter expression, a dose of AAV8-CAG-FLEX-tdTomato was delivered by intracranial (IC) injection into the SN. Unfortunately, neither *Nmur1*^*iCre*^ nor *Cckar*^*iCre*^ mice exhibited detectable tdTomato reporter expression co-localizing with TH staining in the SN (**Figure 3A-B**). To verify the functionality of this AAV, we injected a different GPCR-Cre expressing line, *Neurotensin receptor 1*^*Cre*^ (*Ntsr1*^*Cre*^)^6^ with the same aliquot of virus in the SN and observed robust tdTomato labeling that showed extensive co-localization with TH (n=1, 32% co-localization in SN and 25% in VTA, **Figure 3C**). Together, these results demonstrate that the viral approach is effective and that the lack of labeling observed in *Nmur1*^*iCre*^ and *Cckar*^*iCre*^ mice reflects absence of functional Cre activity rather than technical failure.

**Figure 3.**
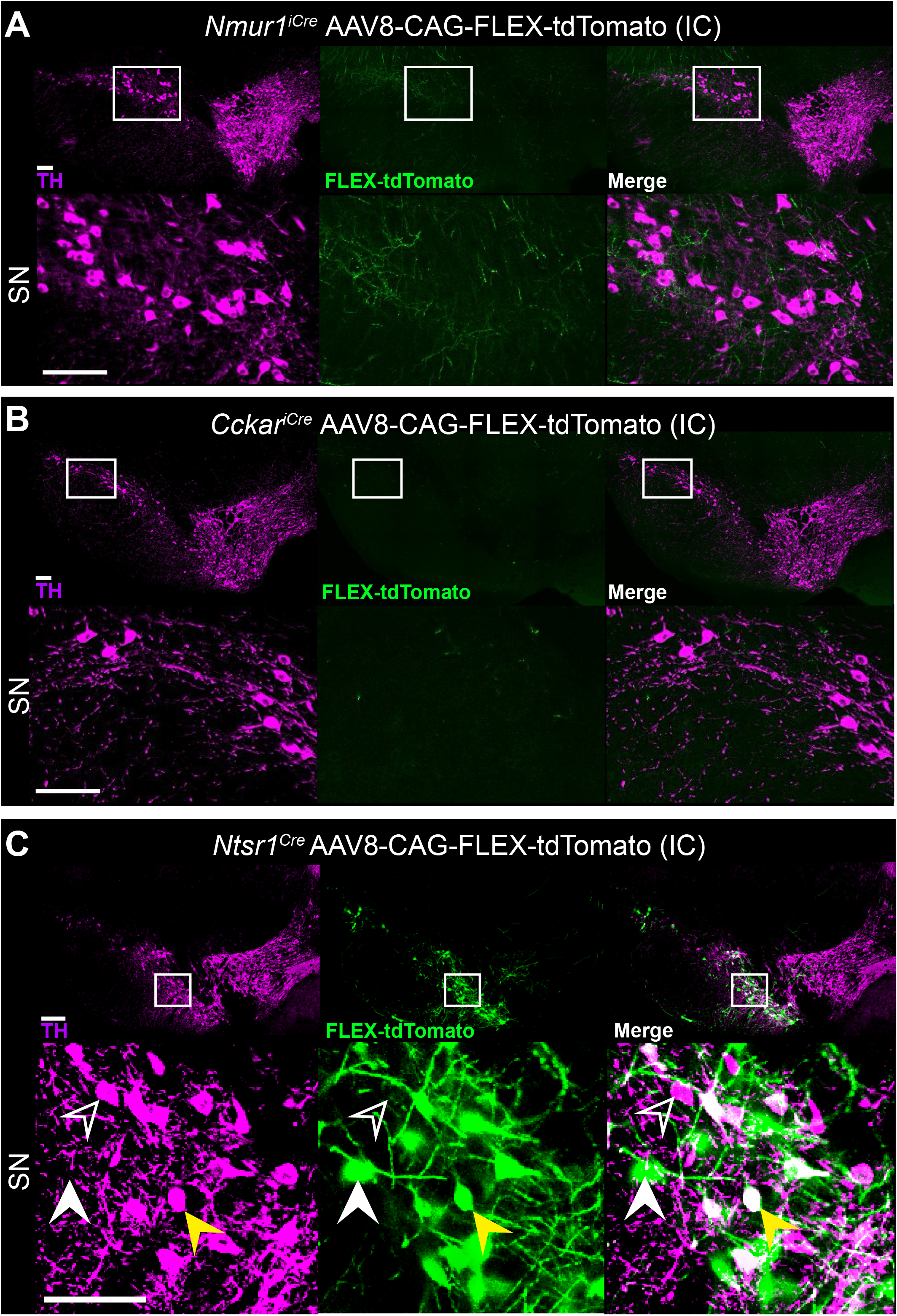
Intra-Cranial injection of AAV8CAG-FLEX-tdTomato expression in the SN and VTA of *Nmur1*^*iCre*^, *Cckar*^*iCre*^, and *Ntsr1*^*Cre*^ mice. (A-C) Representative images of AAV8-CAG-FLEX-tdTomato reporter expression in the SN and VTA from *Nmur1*^*iCre*^, *Cckar*^*iCre*^ and *Ntsr1*^*Cre*^ mice. TH immunostaining (magenta) and FLEX-tdTomato expression (green). Merged image showing colocalization of FLEX-tdTomato and TH (white). Insets show higher-magnification views of boxed regions. Yellow arrowheads indicate TH^+^/FLEX-tdTomato colocalization. Open arrowheads indicate TH^+^ cells. White arrowheads indicate FLEX-tdTomato reporter. Scale bars =100 *µ*m.

Adult recombination was further assessed following retro-orbital (RO) injection of a blood-brain barrier crossing AAV, PHP.eB-CAG-Flex-tdTomato. However, *Nmur1*^*iCre*^ and *Cckar*^*iCre*^ mice again showed an absence of detectable reporter expression in SN and minimal co-localization in VTA (n=2 0% co-localization in SN and 2.6% in VTA, **Figure 4A-B**). In the *Nmur1*^*iCre*^ sections, tdTomato signal was observed in other brain regions, including the medial geniculate complex, indicating successful viral transduction. Similarly, in *Cckar*^*iCre*^ mice, reporter expression was detected outside of the SN and VTA, with labeling visible within the medial mammillary nucleus and sparse signal in the VTA, where only a few neurons showed co-labeling of TH and tdTomato (n=2 mice). In contrast, an RO injection of PHP.eB-CAG-FLEX-tdTomato was given to *Slc6a3*^*Cre*^ mice as a positive control resulting in robust reporter labeling throughout the SN and VTA, as expected for the dopamine transporter (n=3 mice 33.5% co-localization in SN and 25.8% in VTA, **Figure 4C**).

**Figure 4.**
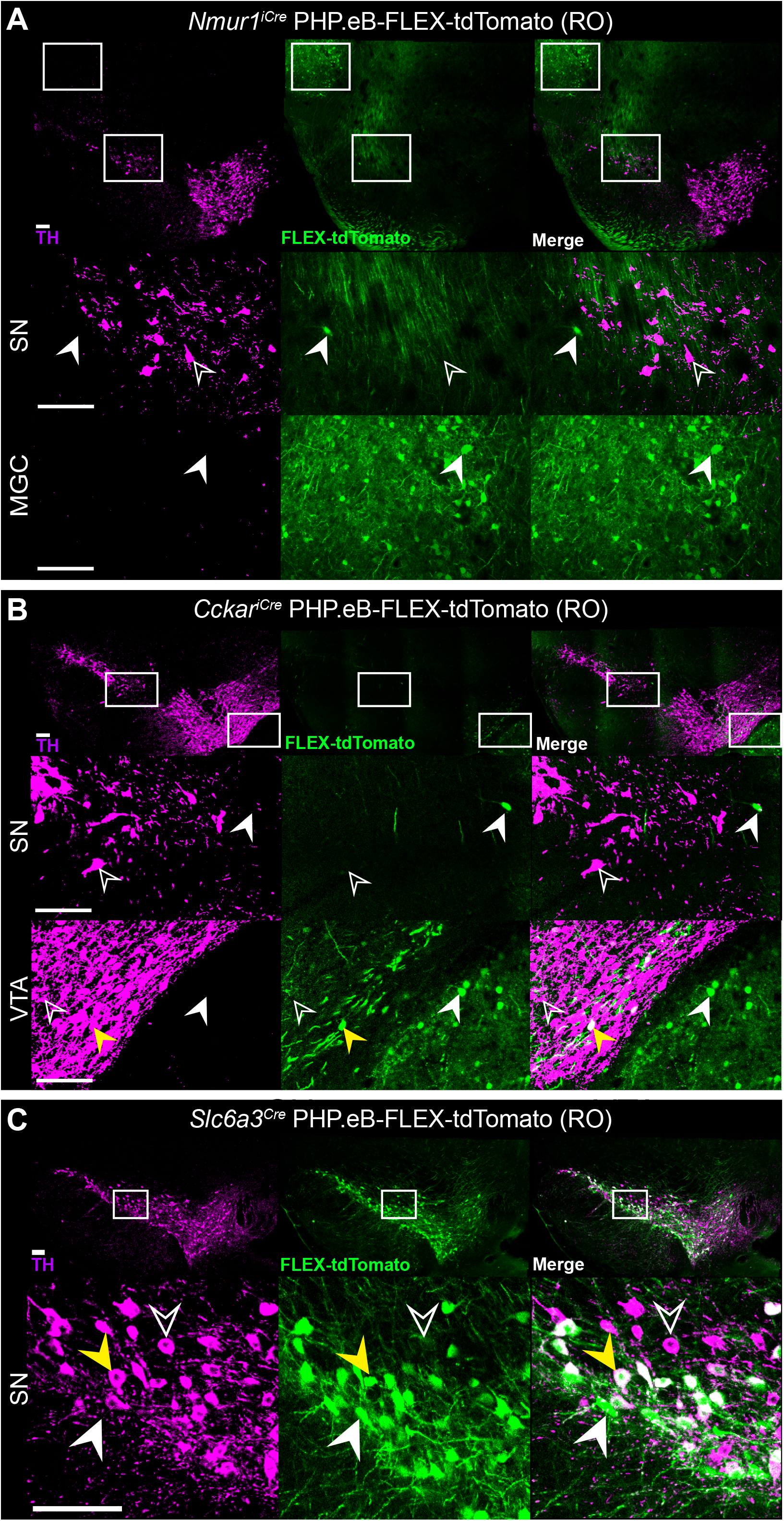
Retro-Orbital injection of PHP.eB-FLEX-tdTomato expression in the SN and VTA of *Nmur1*^*iCre*^, *Cckar*^*iCre*^, and *Slc6a3*^*Cre*^ mice. (A-C) Representative images of PHP.eB-FLEX-tdTomato reporter expression in the SN and VTA from *Nmur1*^*iCre*^, *Cckar*^*iCre*^ and *Ntsr1*^*Cre*^ mice. TH immunostaining (magenta) and FLEX-tdTomato expression (green). Merged image showing colocalization of FLEX-tdTomato and TH (white). Insets show higher-magnification views of boxed regions. Yellow arrowheads indicate TH^+^/FLEX-tdTomato colocalization. Open arrowheads indicate TH^+^ cells. White arrowheads indicate FLEX-tdTomato reporter. Scale bar = 100 *µ*m.

To assess adult transcript localization, we performed hybridization chain reaction (HCR) *in situ* hybridization (ISH) for *Cckar, Th*, and *Dat. Th* and *Dat* signals robustly identified dopaminergic neurons within the adult midbrain. In contrast, *Cckar* signal was observed outside of the *Th*^*+*^ neurons of the SN and VTA, with labeling evident in the medial mammillary nucleus (**Figure 5A**). We next used RNAscope ISH to examine expression of *Ffar4*^4^—the only GPCR atlas candidate reported as validated—alongside *Ntsr1*. We quantified co-localization by sampling 50 *Ntsr1*+ neurons in both the VTA and SN from n = 3 mice and assessing *Ffar4* expression. All *Ntsr1*+ neurons were *Ffar4*+ (150/150 cells). Reciprocal analysis of 50 *Ffar4*+ neurons per region likewise showed complete overlap with *Ntsr1* (150/150 cells, **Figure 5B**). We examined MERFISH data for expression of GPCRs *Ntsr1* and *Crhr1*, determining that they are highly enriched in SN and VTA Th+ populations (**Figure 5C**). However, other candidate markers from the GPCR atlas,^4^ such as *Npy1r, Gpr55*, and *Adrb1* were absent (**Supp. Figure 1A-C**).

**Figure 5.**
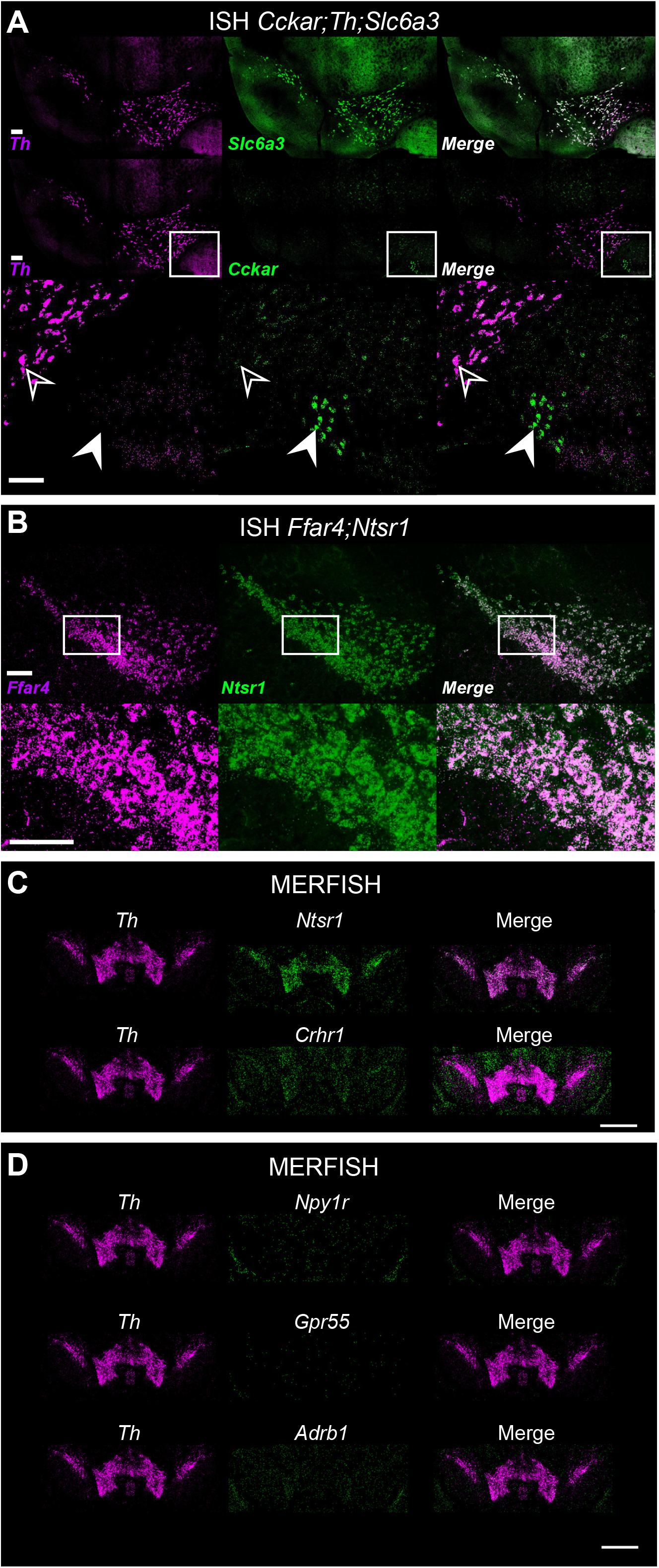
*In situ hybridization* of candidate GPCR marker expression in adult mouse midbrain. (A) Representative images of *in situ hybridization* of *Th* (magenta), *Slc6a3* (green), and *Cckar* (green) in the midbrain of adult mouse tissue. Merged image showing colocalization of *Slc6a3/Cckar* and *Th* (white). Insets show higher-magnification views of boxed regions. Yellow arrowheads indicate *Th*^+^ and *Slc6a3/Cckar* colocalization. Open arrowheads indicate TH^+^ cells. White arrowheads indicate *Slc6a3/Cckar* cells. (B) Representative images of *in situ hybridization* of *Ffar4* (magenta), and *Ntsr1* (green) in the midbrain of adult mouse tissue. Merged image showing colocalization of *Ffar4* and *Ntsr1* (white). Insets show higher-magnification views of boxed regions. Scale bars, 100 *µ*m. (C) Representative images of *MERFISH* of marker genes (*Th, Ntsr1*, and *Crhr1*) and candidate GPCR genes (D) (*Npy1r, Gpr55*, and *Adrb1*). Scale bar = 1 mm.

To further assess the expression of candidate GPCR markers in adult midbrain dopaminergic populations, we examined publicly available spatial transcriptomic datasets, including the Broad Institute^11^ and ABC Atlas,^12^ comparing transcript localization across receptors identified in the GPCR atlas (**Figure 6**). As expected, *Th* signal was robustly detected throughout the SN and VTA in both datasets, confirming reliable identification of dopaminergic neuron populations (**Figure 6A-B)**. In contrast, several candidate receptors, including *Npy1r, Htr5b, Cckar, Npsr1, Mc4r*, and *Npy5r*, exhibited only sparse labeling distributed broadly across the section, without enrichment in dopaminergic midbrain regions (**Figure 6C-N**). *Nmur1* transcripts were largely undetectable in the SN and VTA in the Broad dataset and showed no clear overlap with Th+ domains (**Figure 6O**). Notably, *Ffar4* transcripts were also sparse and not detectably enriched within the midbrain in the Broad dataset (**Figure 6P**), which conflicts with our RNAscope data (**Figure 5B**) and the data of Apuschkin and colleagues (2024).^4^

**Figure 6.**
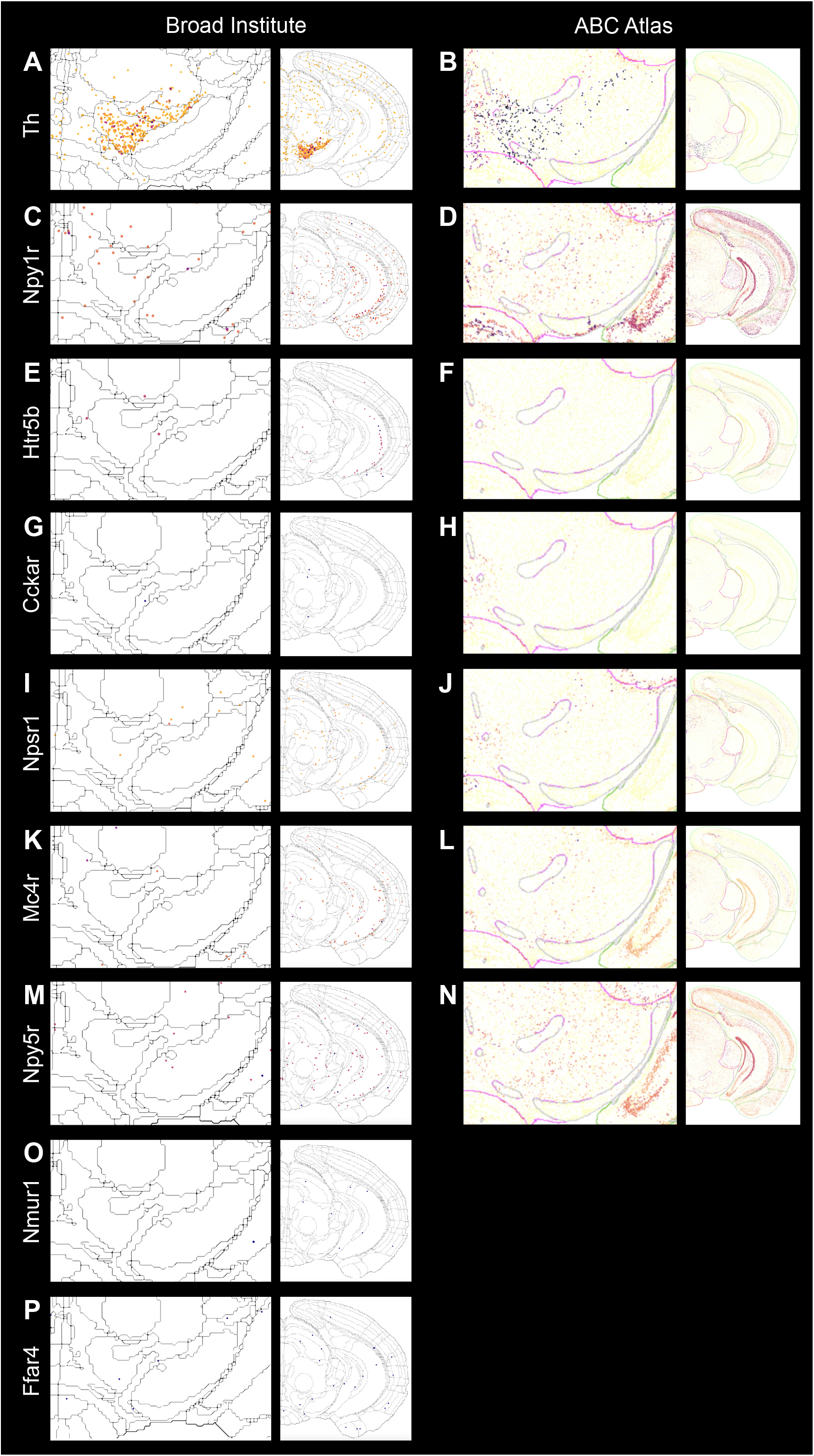
Spatial transcriptomics reveal minimal candidate GPCR marker expression in adult midbrain dopaminergic regions. (A-B) Spatial transcriptomic maps from Broad Institute and Allen Brain Cell (ABC) Atlas showing robust *Th* transcript localization throughout the substantia nigra (SN) and ventral tegmental area (VTA), confirming identification of dopaminergic neuron populations. (C-P) Spatial transcriptomic maps for additional candidate GPCR markers (*Npy1r, Htr5b, Cckar, Npsr1, Mc4r, Npy5r, Nmur1*, and *Ffar4)*.

## Discussion

Our findings demonstrate that *Nmur1* and *Cckar*, identified as candidate GPCR markers of DA neuron subpopulations,^4^ do not reliably reflect adult expression in midbrain DA neurons. Across complementary approaches, including reporter lines, conditional genetic manipulation, viral reporters, spatial transcriptomics, and in situ hybridization, we find no evidence supporting functional expression of *Nmur1* in adult DA neurons. Although Apuschkin et al. (2024) reported reduced *Nmur1* expression in adult DA neurons relative to those isolated early in life,^4^ we were nevertheless surprised to observe little evidence of developmental *Nmur1*^*iCre*^ activity, as reflected by the lack of *Th* deletion and the minimal Ai6 reporter expression in midbrain DA neurons (2% in SN and 5% in VTA). In contrast, *Cckar* exhibits a clearer dissociation between minimal or undetectable adult transcript expression and measurable recombination as assessed by both *Th* deletion and Ai6 reporter activity, consistent with developmental or transient activity rather than stable adult identity.

These results highlight an important limitation in the interpretation of transcriptomic enrichment as a proxy for functional marker validity, particularly when extrapolating to adult neuronal identity. While prior work has demonstrated that transcriptomic datasets can successfully define biologically meaningful dopaminergic subpopulations,^3,13,14^ these studies have relied on convergence between molecular identity, spatial localization, and functional validation. For example, recent large-scale transcriptomic and spatial mapping efforts have resolved midbrain DA neurons into organized populations with reproducible anatomical and functional correlates.^3,14^ In contrast, the GPCR signals identified here do not consistently generalize to adult expression when evaluated using independent experimental approaches.

To further assess the generalizability of these findings, we examined publicly available spatial transcriptomic datasets of the adult mouse brain, including the Allen Brain Cell Atlas and Broad Institute resources, as well as our own MERFISH analyses. Across these independent datasets, we observe limited or absent expression of many of the candidate GPCRs within dopaminergic neuron populations, including *Nmur1* and *Cckar*. These observations are consistent across platforms and analysis approaches, suggesting that the discrepancies identified here are not restricted to a single experimental modality. An exception is *Ffar4*, which was detected in both the original study^4^ and in our RNAscope analyses but shows minimal or no enrichment within dopaminergic midbrain regions in the Broad spatial transcriptomic datasets, highlighting platform-dependent differences in sensitivity and detection. Additional inconsistencies between transcriptomic predictions from Apuschkin et al. (2024) and established observations further underscore these limitations. For example, their reported low prevalence of *Ntsr1* expression in DA neurons contrasts with numerous published studies,^6,7,15^ MERFISH data presented here, and our observations using a Cre-dependent AAV reporter, all of which support broader representation within midbrain dopaminergic populations. Moreover, the only validated candidate GPCR from the atlas, *Ffar4*, appears to substantially overlap with the well-characterized *Ntsr1* population. Pharmacological manipulation of *Ffar4* was reported to influence feeding and drinking behaviors,^4^ which are phenotypes previously associated with *Ntsr1*-expressing DA neurons.^16–19^ These observations raise the possibility that *Ffar4* does not define a distinct DA class but instead reflects a molecular signature of the already established and well-characterized *Ntsr1* population. GPCRs previously implicated in DA neuron function, including *Crhr1*, are not represented in the GPCR atlas.^20^ While differences in datasets and detection approaches may contribute to these discrepancies, they collectively highlight that transcriptomic datasets alone may not fully capture functionally relevant receptor expression.

Importantly, these conclusions do not negate the utility of transcriptomic atlases for identifying candidate genes but instead emphasize the need for careful validation prior to their use in genetic targeting strategies. This distinction is particularly relevant for GPCRs, which are frequently prioritized for their pharmacological and experimental accessibility and for which Cre driver lines are often readily available. As a result, transcriptomic enrichment can be rapidly translated into experimental targeting approaches. Without validation in adult tissue, however, these markers can be of limited utility for circuit-specific manipulations occurring in adults. We propose that candidate markers emerging from transcriptomic datasets should be evaluated using a minimal validation that includes: (1) assessment of co-localization with established cell-type markers in adult tissue, (2) confirmation of Cre-dependent recombination in adult animals, and (3) comparison across independent datasets where possible. Incorporating such validation pipelines alongside transcriptomic discovery efforts will improve the reliability of candidate markers and facilitate their translation into experimental applications.

## Methods

### Experimental Model and Subject Details

The experiments described herein were approved by the California State Polytechnic, Pomona Institutional Animal Care and Use Committee (IACUC) under protocols: 23.006, 23.009, 25.018.

Mice were group housed under a 12 h light/dark cycle with *ad libitum* access to food and water. Housing conditions were maintained at an ambient temperature of 21-24 °C and relative humidity between 20–45%. Cre driver lines for *Nmur1* and *Cckar* were obtained from The Jackson Laboratory. Cre lines were crossed to the *Ai6* Cre-dependent reporter line (*Rosa26-loxP-STOP-loxP-ZsGreen*; The Jackson Laboratory). Both male and female mice were used for experiments. Adult mice (≥8 weeks of age) were used for all analyses.

Genomic DNA was obtained from tail biopsies collected from mice at 12–14 days of age. Samples were submitted to Transnetyx (Cordova, TN) for analysis. Genotyping was performed using real-time PCR to detect the presence of *iCre* or *Cre* recombinase, *Th flox* allele, *wild-type Th* allele, and the *ZsGreen* allele.

### Tissue preparation and immunohistochemistry

Transcardiac perfusion was performed by injecting 5-10 mL of 0.1 M phosphate buffer solution into the left ventricle followed by 5 mL of 4% PFA (Fisher Scientific), both made fresh prior to perfusion. Whole brain tissue was removed and stored in 4% PFA at 4°C for 24-hours, then placed in 0.1 M PBS. 50-*µ*m coronal sections were obtained using a Leica VT1000S Vibratome (Leica Biosystems, Buffalo Grove, IL) and stored in 0.1M PBS at −4°C. For staining, tissue samples were placed in glass spot plates in 0.1 M PBS for 1-min, then permeabilized by incubating in PBST (0.5% Triton X-100 in PBS) for 10-min. Samples were then blocked in 5% goat serum (Gibco, #16210-064) solution for 10-min. Tissue samples were then transferred to a well of 1:500 diluted primary antibody, Aves chicken anti-TH (Antibodies, Inc) diluted in goat serum overnight at 4°C. After primary antibody staining, tissue was then washed three times in PBS for 5-min each. Tissue was then exposed to a 1:500 dilution of Goat anti-chicken Alexafluor 647 secondary antibody (Thermofisher) in PBS for 1-hour shielded from light at room temp. The tissue samples were washed in PBS three times for 5-min each before mounting using Epredia Immu-Mount (FisherScientific, #9990402) and cover slipping.

### Fluorescence imaging and analysis

Images were acquired using a Nikon C2 Confocal scope with a standard detector system and Nikon Elements Software with identical acquisition settings across experimental groups. Co-localization between Cre-dependent reporter expression (ZsGreen or tdTomato) and TH immunoreactivity was assessed manually or using ImageJ/Fiji. Cells were considered co-localized if reporter signal overlapped with TH signal within the same soma. Quantification was performed on multiple sections per animal across rostrocaudal levels of the midbrain. Data are presented as mean ± SEM.

### In situ hybridization

Multiplex fluorescent *in situ* hybridization was performed using the HCR *in situ* hybridization (Molecular Instruments) according to the manufacturer’s instructions. Probes targeting *Cckar, Th*, and *Slc6a3* (*Dat*) were used. Sections were imaged using confocal microscopy, and co-localization was assessed by identifying cells containing punctate signal for each probe within the same soma. Quantification was performed across multiple sections and animals. RNAscope for *Ntsr1* and *Ffar4* were performed as previously described.^21^ Quantification of RNAscope data was performed across multiple sections obtained from n=3 mice. For each section, cells were counted in one fluorescence channel, using a synchronized cell counter (FIJI), and corresponding fluorescence channel was assessed to determine co-localization. This procedure was then repeated to the corresponding fluorescence channel.

### Multiplexed error-robust fluorescence in situ hybridization (MERFISH) and data analysis

MERFISH assays were conducted as previously described.^22^ Briefly, fresh-frozen coronal brain sections (12 μm thick) encompassing the SN and VTA from the respective mice were prepared using a Leica CM1860 cryostat and processed according to the Vizgen MERSCOPE Sample Preparation User Guide. A custom-designed mouse gene panel (Vizgen #VZG0191) was used for hybridization. The panel consisted of binary-coded probes targeting 500 selected genes, including markers for various brain cell types and genes involved in multiple signal transduction pathways. Hybridization was performed at 37 °C for approximately 40 h, followed by sequential washes in formamide buffer (30 min at 47 °C, twice), gel embedding, and protease K–mediated tissue clearing overnight at 37 °C. Imaging was performed on the Vizgen MERSCOPE Ultra using the manufacturer’s 500-gene kit after assembly of the cleared slides into the FCX-S gasket for image acquisition. Image acquisition was programmed using Vizgen MERSCOPE software (version 234c.250721.1718) under default 3D settings with both polyT and DAPI channels enabled and a scan depth of 10 μm. Upon completion of MERFISH imaging, the generated files were exported for downstream analysis using MERFISH Visualizer (version 2.5.3504.0) and custom-written analysis code.

MERFISH datasets from two mice samples were analyzed to characterize GPCR expression within dopaminergic brain regions. Cells were assigned to manually annotated SN and VTA regions based on spatial coordinates derived from the imaging dataset, and only cells located within these regions were included in downstream analyses. Single-cell transcript counts for *Th* and 13 GPCR genes were quantified after removal of low-confidence transcripts (transcript score ≤ 0.7) and invalid cell assignments. Cells were classified as gene-positive when more than two transcripts were detected for a given gene. The abundance of *Th*/*GPCR* double-positive cells was then quantified across samples and anatomical regions.

### Conditional genetic manipulation

Conditional knockout mouse lines were generated using Cre-loxP recombination targeting *Th*. The following lines were used: *Cckar*^*em1(icre)Shah*^*/J* (037017), *Slc6a3*^*tm1(cre)Xz*^*/J* (020080), *Nmur1*^*iCre-eGFP*^ (038197), *Gt(ROSA)26Sor*^*tm6(CAG-ZsGreen1)Hze*^ (007096), and *Ntsr1*^*Cre*^ (033365). All strains were obtained from The Jackson Laboratory (strain numbers indicated in parenthesis except for the *Ntsr1*^*Cre*^ mice, which were obtained from the breeding colony of Dr. Gina Leinninger of Michigan State University).

### AAV production

Recombinant AAVs were produced and purified according to the protocol described by Challis et al. (2019).^23^ HEK293T cells were transfected with a triple-plasmid system consisting of a capsid (pUCmini-iCAP-PHP.eB, Addgene #103005 or pAAV2/8, Addgene #112864), cargo (AAV pCAG-FLEX-tdTomato-WPRE, Addgene #51503), and pHelper (NovoPro #V005569) plasmids using polyethylenimine. Viral particles were harvested from culture media and cells before purification via ultracentrifugation in a discontinuous iodixanol density gradient. Viral particles were then washed and concentrated via centrifugal ultrafiltration with concomitant buffer exchanges using Dulbecco’s Phosphate Buffered Saline (DPBS) containing 0.001% Pluronic F-68 to prevent viral aggregation and minimize adherence to plasticware. Purified AAVs were titered (vg/*µ*L) via digital PCR using the QuantStudio Absolute Q Digital PCR System utilizing probes targeting inverted terminal repeats.

### Systemic AAV delivery

For systemic viral delivery, the Cre-dependent reporter PHP.eB-CAG-FLEX-tdTomato was administered to adult mice via retro-orbital injection. Mice were anesthetized with isoflurane (5% induction, 1–3% maintenance in 1.5 L/min O_2_) and positioned prone. A 50 *µ*L volume containing 3 × 10^11^ viral genomes (vg) was injected into the right retro-orbital sinus using a sterile 31-gauge insulin syringe.

### Stereotaxic Intracranial AAV Delivery

Stereotaxic delivery of AAV serotype 8 was used to assess Cre-dependent recombination in *Nmur1*^*iCre*^, *Cckar*^*iCre*^, and *Ntsr1*^*Cre*^ mice. AAV8-CAG-FLEX-tdTomato was injected as a Cre-dependent reporter at a dose of 4.0 × 10^8^ vg per injection site.

Mice underwent surgery at 8–12 weeks of age. Mice undergoing surgery were first induced in a sealed induction chamber with an oxygen flow rate of 1 lpm OL and 5% isoflurane. Once fully anesthetized, mice were placed onto a stereotaxic frame (Kopf, Model 1900 Stereotaxic Alignment System) and stabilized. The scalp was sterilized with three washes of chlorhexidine followed by isopropanol, and an incision was made to expose the skull. Excess fascia was removed with sterile cotton swabs to allow for full exposure of the skull. Once exposed, the skull was leveled (Kopf, Model 1905 Stereotaxic Alignment Indicator), and the drill was centered on bregma, the intersection of the coronal and sagittal sutures of the skull. From this point, holes were drilled in the skull using coordinates to specifically target the substantia nigra (x = ±1.5 mm, y = ™3.6 mm). A microinjector with an attached glass pipet loaded with virus was slowly lowered into the brain to the proper depth (™4.0 mm). After waiting 5 min, bilateral injections of 750 nL of virus were given at a delivery rate of 10 nL/second, followed by a 10-min waiting period following each injection to optimize uptake of virus into surrounding tissue. Following injection, the skull was sealed with bone wax (Surgical Specialties, #309) and 0.1 mL topical anesthetic (Hospira, Marcaine 0.5%, 5 mg/mL, PAA113011) was applied to the skull and incision site prior to sealing the incision with VetBond (3M, No. 1469SB). After isoflurane was turned off, an additional 0.5 mL sterile saline was injected subcutaneously. Mice were transferred into a sterile home cage placed beneath a heating lamp until consciousness returned. Animals were weighed before and after the procedure to account for weight gain due to supplementary fluids. Animals were monitored for a minimum of three days following surgery and were given 0.1 mL 0.5 mg/mL ketoprofen and Nutri-Cal (Vetoquinol) daily; an additional 0.5 mL saline SQ could also be provided at researcher’s discretion, based on animal condition.

### Statistical analysis

Statistical analyses were performed using GraphPad Prism 11.0. Data are presented as mean ± SEM. Comparisons between groups were made using a t-test. Statistical significance was defined as p < 0.05.

## Acknowledgment

Funding was provided by the National Institute of General Medical Sciences of the National Institute of Health under award number R16GM145576, National Institute of Diabetes, Digestive and Kidney Disorder under award number DK-132736, and BRAIN Initiative Armamentarium for Precision Brain Cell Access U24MH131054 to ADS. The content is solely the responsibility of the authors and does not necessarily represent the official views of the National Institutes of Health. MS is a recipient of funding from a T32GM137812 National Institute of General Medical Sciences award at Cal Poly Pomona.

## Supplemental Figure Legend

**Figure S1. MERFISH data analysis of candidate GPCR expression in adult mouse midbrain**. (A) Representative images of MERFISH data analysis of marker genes (*Th, Ntsr1*, and *Crhr1*) and candidate *GPCR* genes (*Npy1r, Gpr55*, and *Adrb1*). (B) Higher-magnification views of MERFISH data analysis of marker genes and candidate GPCR genes. (C) Percentage of *Th*^+^ cells that are *GPCR*^+^ in the dopaminergic midbrain region. Data are represented as mean ±SEM. n = 2 mice, 4 samples for each mouse. Scale bars = 100 *µ*m.

### Declaration of generative AI and AI-assisted technologies in the manuscript preparation process

During the preparation of this manuscript, the authors used ChatGPT and Gemini to improve the clarity and readability of the text. After using this tool/service, the authors reviewed and edited the content as needed and take full responsibility for the content of the published article.

## Notes

### Competing Interest Statement

The authors have declared no competing interest.

