## Supplementary figures and images for "Nmur1 and Cckar fail to support functional genetic access in adult dopamine neurons and challenge GPCR atlas assignments"

### Supplemental Figure 1

#### Supplemental 1

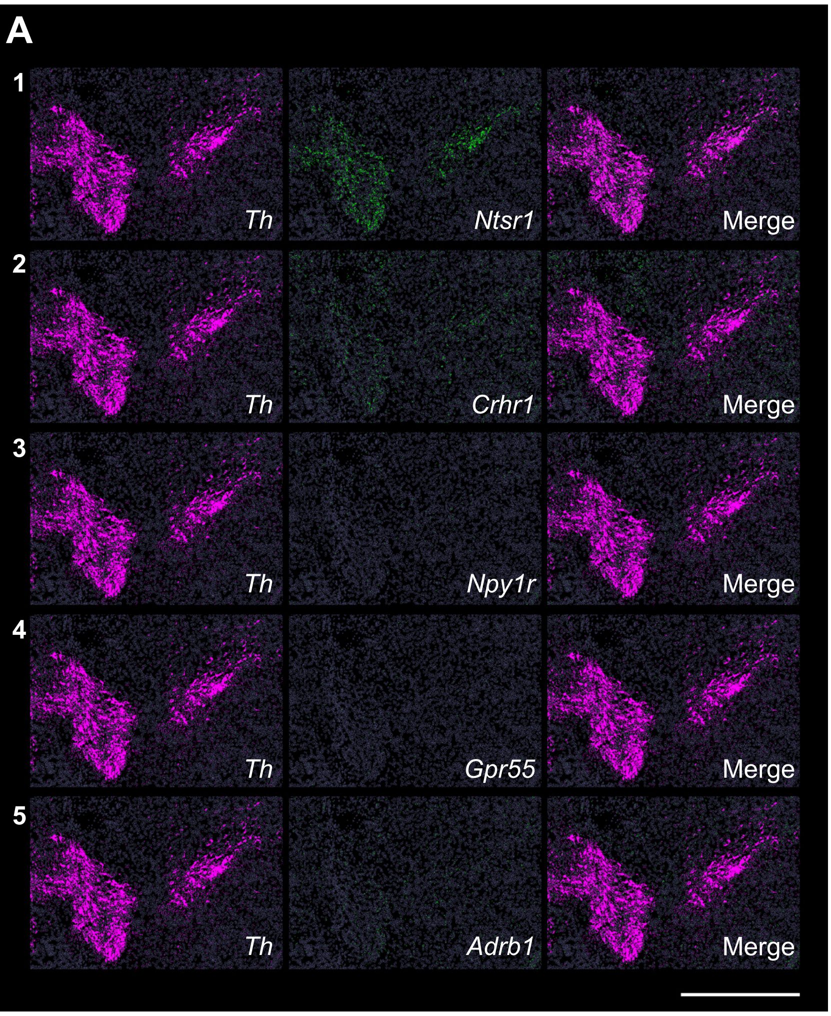


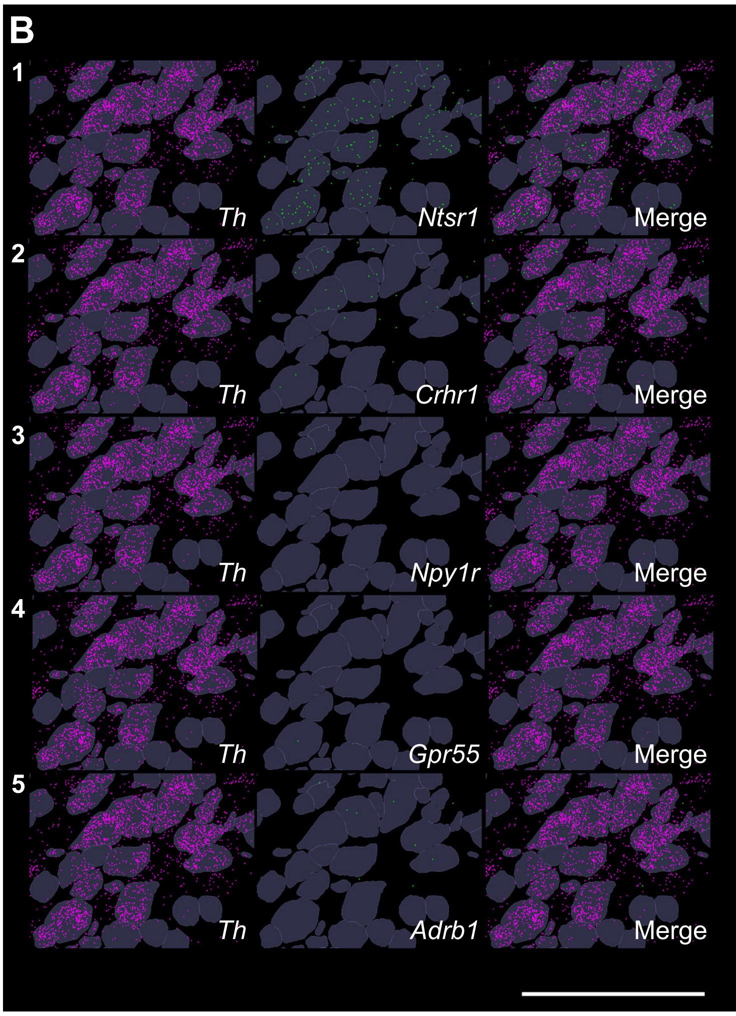


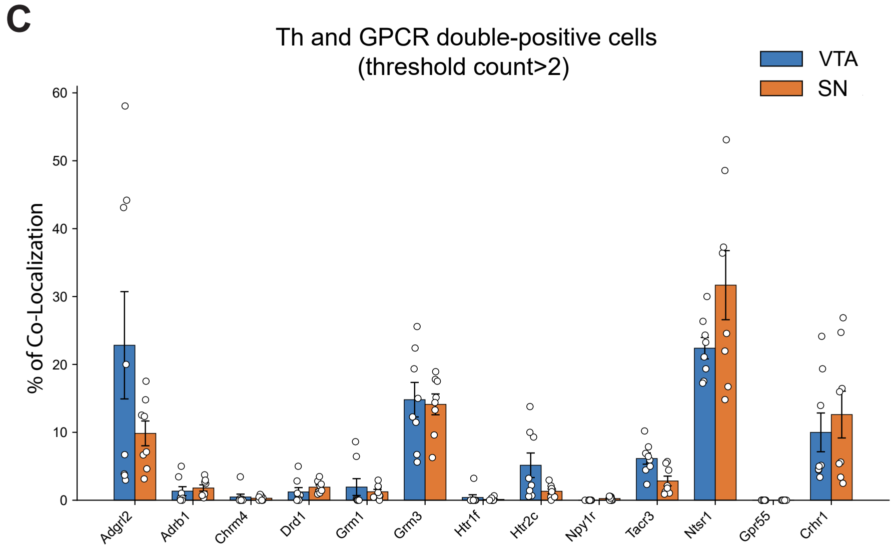
